# Beyond linearity in neuroimaging: Capturing nonlinear relationships with application to longitudinal studies

**DOI:** 10.1101/2020.11.01.363838

**Authors:** Gang Chen, Tiffany A. Nash, Katherine M. Reding, Philip D. Kohn, Shau-Ming Wei, Michael D. Gregory, Daniel P. Eisenberg, Robert W. Cox, Karen F. Berman, J. Shane Kippenhan

**Author notes:** These two authors contributed equally (Karen F. Berman, J. Shane Kippenhan).

## Abstract

The ubiquitous adoption of linearity for quantitative explanatory variables in statistical modeling is likely attributable to its advantages of straightforward interpretation and computational feasibility. The linearity assumption may be a reasonable approximation especially when the variable is confined within a narrow range, but it can be problematic when the variable’s effect is non-monotonic or complex. Furthermore, visualization and model assessment of a linear fit are usually omitted because of challenges at the whole brain level in neuroimaging. By adopting a principle of learning from the data in the presence of uncertainty to resolve the problematic aspects of conventional polynomial fitting, we introduce a flexible and adaptive approach of *multilevel smoothing splines* (MSS) to capture any nonlinearity of a quantitative explanatory variable for population-level neuroimaging data analysis. With no prior knowledge regarding the underlying relationship other than a parsimonious assumption about the extent of smoothness (e.g., no sharp corners), we express the unknown relationship with a sufficient number of smoothing splines and use the data to adaptively determine the specifics of the nonlinearity. In addition to introducing the theoretical framework of MSS as an efficient approach with a counterbalance between flexibility and stability, we strive to (a) lay out the specific schemes for population-level nonlinear analyses that may involve task (e.g., contrasting conditions) and subject-grouping (e.g., patients vs controls) factors; (b) provide modeling accommodations to adaptively reveal, estimate and compare any nonlinear effects of an explanatory variable across the brain, or to more accurately account for the effects (including nonlinear effects) of a quantitative confound; (c) offer the associated program **3dMSS** to the neuroimaging community for whole-brain voxel-wise analysis as part of the AFNI suite; and (d) demonstrate the modeling approach and visualization processes with a longitudinal dataset of structural MRI scans.

## 1 Introduction

There are two types of variables in statistical modeling. The first type is categorical (e.g., sex, group, condition), and its possible categories are treated as the levels of a factor. The other type is quantitative, in which numerical values represent measurements (e.g, BOLD response, age, reaction time). Sometimes a variable can be treated one way or another depending on the research focus and hypothesis. For example, when a group of subjects are scanned once during each of five consecutive years, the five time points can be modeled as a factor with 5 levels when the differences among them, irrespective of order, are of interest. On the other hand, if the investigator wants to probe the trend over time, the same five points can be modeled as values of a quantitative variable, typically some version of the subject’s age.

From a temporal perspective, there are two types of experimental study designs: cross-sectional and longitudinal. A cross-sectional study compares single-time-point observations or measurements across different groups of subjects without respect to time. In contrast, a longitudinal study involves repeated observations or measurements of the same subjects over a defined period of time, with potentially variable times for subjects’ individual observations. Longitudinal studies are of great import in neuroimaging, addressing fundamental issues such as neurodevelopment, aging, and medication effects over time. An important benefit of a longitudinal study is that the investigator can detect changes or developmental trajectories at both the population and subject level, extending beyond a single point in time and potentially providing greater opportunity to assess causal contributors to dependent variables. In general, the choice between cross-sectional and longitudinal designs will be driven by the nature of the research goal and associated issues of practicality. For example, an investigator interested in understanding group differences in a measurement (e.g., between individuals with and without an illness) will be well-served by a cross-sectional design, whereas if the hypothesis being tested involves changes over time (e.g., the trajectory of a measurement over the course of an illness’ progression), a longitudinal design may be more appropriate. However, some temporal questions may be better suited to cross-sectional study because of pragmatic considerations, such as time constraints, as generally, cross-sectional studies can be executed more quickly than longitudinal studies. An investigator studying how a measurement changes across the entire human lifespan, for instance, may select a cross-sectional design so that the study can be completed within their lifetime.

The proper handling of a quantitative explanatory variable (especially in a longitudinal dataset) can be a daunting task for an analyst or modeler. When such a variable is incorporated into a model, the investigator may be interested in either exploring its effect or controlling for its variability. A typical approach is to assume a linear relationship. From the modeling perspective, a longitudinal study is signified by its specific treatment of the time variable and the dependence of repeated measurements collected. Depending on the number of time points, the analyst may treat the time variables as categorical or continuous/quantitative. For instance, when only a few time points are involved or the order of the time points is not critical, one may simply consider them as the levels of a within-subject or repeated-measures factor. The general linear model (GLM) is a powerful statistical tool especially when no within-subject or repeated-measure factors (e.g, a task factor with three levels of positive, negative and neutral) are involved. However, despite the relative simplicity, correctly incorporating a repeated-measures factor into a population-level model through a univariate GLM framework remains a challenge in neuroimaging (Chen et al., 2014; Telzer et al., 2018; McFarquhar, 2019), even though more flexible and appropriate frameworks such as multivariate GLM and linear mixed-effects (LME) modeling have been used for decades. Specifically, whenever a repeated-measures factor is involved, the univariate GLM framework may struggle to properly partition the relevant effects due to the difficulty of accurately characterizing the multiple levels embedded in the data hierarchy and can be further hamstrung by its inability to handle the presence of any quantitative explanatory variables (Chen et al., 2014). These limitations can be readily addressed under a multivariate GLM framework (e.g., as implemented in the program **3dMVM** in AFNI). An additional consideration is that missing data are very common in longitudinal studies, presenting another challenge for population-level analysis, and the LME platform (Chen et al., 2013) can effectively characterize the data variability through the use of variance-covariance structures such as varying intercept/slope and stratified/crossed effects (e.g., as implemented in the programs **3dLME** and **3dLMEr** in AFNI) and can handle missing data as long as the absences can be considered random.

One may convert a quantitative explanatory variable into a factor by categorizing the quantitative variable into two or more intervals for a conventional ANOVA. However, this methodology of binning or discretization should be discouraged despite its expediency. First, it can lead to the loss of information, precision and inferential power. Any arbitrariness in the choice of cutoff points comes with an assumption of equal intervals between consecutive bins and artificial discontinuities at the cutoff points. Representing the variables as continuous will avoid information loss, but the default approach when handling the effect of a quantitative variable is to assume linearity. With rare exceptions, linearity is the underlying assumption for modeling a quantitative covariate, including, to some extent, approaches using a higher-order relationship (e.g., quadratic or cubic), since these can be considered as special cases of interactions (i.e., a variable interacting with itself).

Most statistical models are constructed with an assumption of linearity for the sake of simplicity. A linear relationship has two properties per the superposition principle: additivity (i.e., aggregate responses reflect the sum of individual component effects) and homogeneity of degree one (i.e., scaling effects by a given factor results in an equivalently scaled response). These qualities make linearly parametric models fairly intuitive to construct, which has likely contributed to their widespread adoption in the literature. In addition, solving a linear system is relatively economic computationally, allowing solutions within a reasonable period of runtime. Linearity might pragmatically be a reasonable approximation of the first-order Taylor expansion^1^ in some cases, especially when the range of the explanatory variable is relatively small (e.g., age spanning over a few years). However, even if an explanatory variable can be mechanically treated as a linear effect in the model, one cannot expect that the linearity assumption would be always reasonable. For example, a simple variable such as reaction time may have a nonlinear effect in some regions of the brain (Chen et al., 2020b). Similarly, processes of human brain development or aging (e.g., measured functionally with regional BOLD response or structurally with anatomical scans) are not necessarily expected to follow a strictly linear trajectory across different life stages (e.g., Fjell et al., 2013; Faghiri et al., 2019).

To account for nonlinear relationships, one may increase the order of polynomials from linearity to a higher order. Polynomial models are popular for several reasons, including their simple formulation, well known and easily-understood properties, relatively flexible shapes, and low computational cost. However, such a strategy faces several challenges.

- The selection of the order of the polynomials can be complicated and arbitrary. It is also difficult to predetermine the order of polynomial fitting, especially with the heterogeneity across brain regions: one particular order of polynomials may work for some regions but not necessarily for others.
- One may have a poor trade-off between model complexity and goodness of fit. For instance, a lower order (e.g., quadratic) polynomial might not be flexible enough to account for adequate variance, while a higher order curve could track the data too closely (leading to potential overfitting), which could cause numerical stability problems (Wood, 2017) or incur artificial oscillations at the edges of an interval over a set of equally-spaced interpolation points (e.g., Runge’s phenomenon (Runge, 1901)).
- It is difficult to assess the statistical evidence for the overall difference between two curves. Even if one could identify the specific terms (e.g., linear, quadratic or cubic) with strong evidence, the interpretation tends to be unwieldy and murky when one addresses such a question as what a cubic term means. For this reason, fitting a polynomial essentially amounts to imposing a predetermined and likely unverifiable structure on the data, rather than conforming the relationship to the data.
- Non-locality or instability is an undesirable property with polynomial modeling: the fitted curve at a particular location may be sensitive to the data far from that point. For example, the twisted turn of the fitted cubic polynomials on the upper left corner in Fig. 1a occurs because of the steep drop at the lower right corner.
- It is common to see polynomials perform poorly in interpolatory, extrapolatory and asymptotic settings, as documented in the literature (Gelman and Imbens, 2019). For example, even if polynomials fit well within the data at hand, their performance will usually deteriorate rapidly outside the data range; in fact, the two ends of any polynomial fit will extend to —∞ and +∞, respectively.

**Figure 1:**
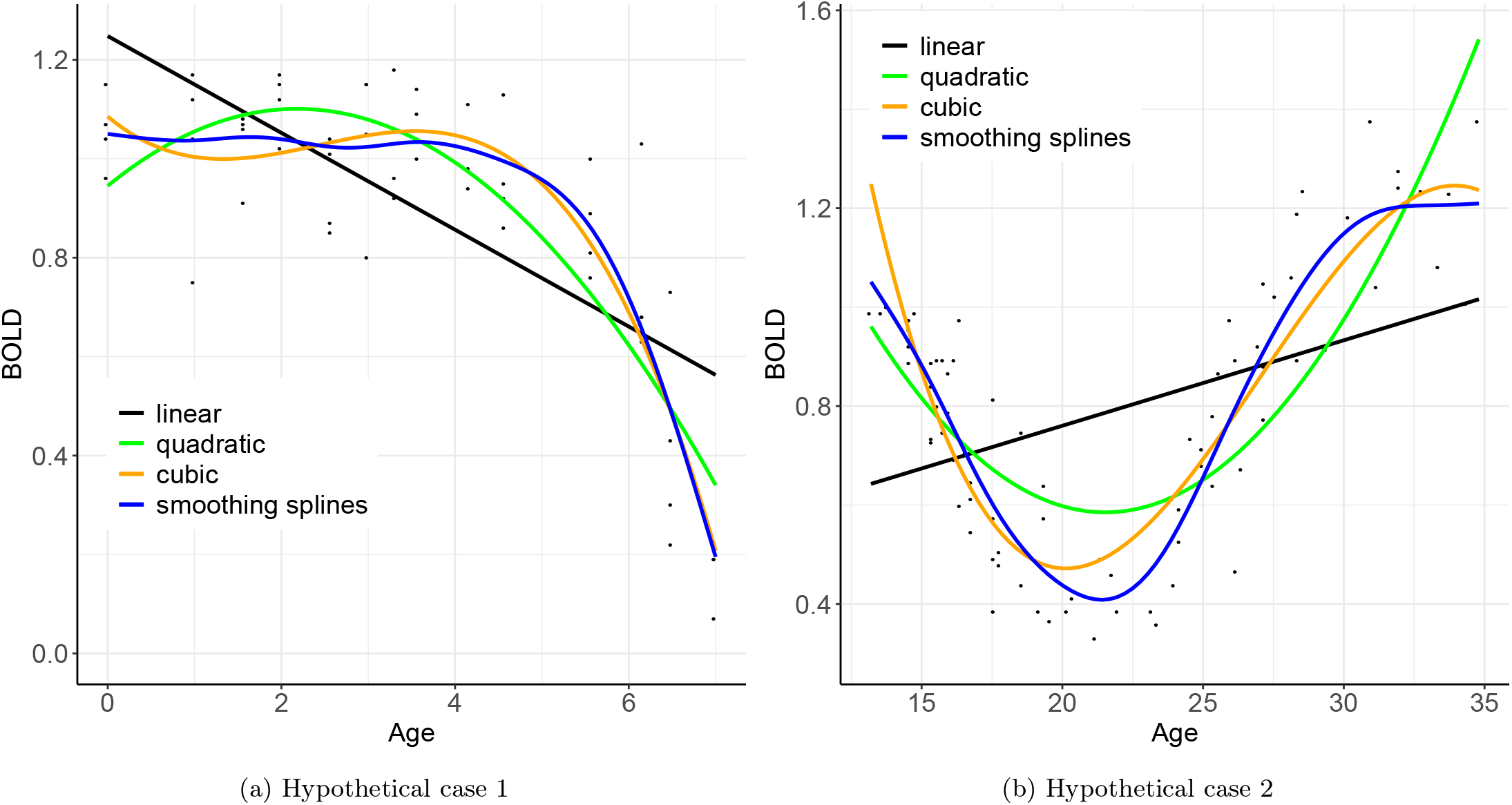
Comparisons of data fitting among three polynomial models and a smoothing spline approach. Linear fitting (black line) between BOLD response and age in a region, as typically adopted in neuroimaging analysis, sometimes may render a roughly acceptable performance with a general sign (positive or negative) for the association between *x* and *y* such as (a) here, but other times it could fail badly as shown in (b). Quadratic fitting (green) performs better, and the cubic model (orange) is a further improvement. Furthermore, the curve estimated through smoothing splines (blue) provides the best fit without any prior knowledge of parameters such as the order of polynomials. More crucially, smoothing spline modeling is largely self-adaptive without the daunting undertaking of order selection in polynomial fitting.

Modeling adaptivity is crucial in neuroimaging because the field faces unique challenges in model formulation, selection and improvement due to the large amounts of data involved. With the massively univariate approach, which is very common in neuroimaging, the same model formulation is applied across the whole brain with varying response variable values. As a result, the typical adoption of a linear fit approach for a quantitative variable of interest or no interest, could be hit-or-miss, as illustrated in Fig. 1. While a linear fit might be reasonable in one region, a cubic model could be required for another region. The flexibility of polynomial fitting, model selection and improvement is basically impeded by imposing the same polynomial order or model formulation across the brain, an organ whose development and functional biology are characterized by profound regional diversity. Further, because of these impediments, model assessments and fit visualizations are cumbersome with whole-brain data. Besides polynomial fitting, another possibility is to adopt a specific nonlinear function or transformation such as exponential or logarithmic relationships. However, this option usually requires some prior knowledge about the underlying mechanism of the trajectory. When such a prerequisite is absent, which is generally the case, we are left with the demand for a more flexible approach to handling nonlinearity.

Here we describe and propose a novel approach to modeling a quantitative explanatory variable in neuroimaging data analysis: multilevel smoothing spline (MSS). Essentially, with MSS, the analyst does not pre-establish a specific functional form relating the variables; rather, it is the data that determine the shape and specifics of the relationship. This methodology, as an alternative to the ubiquitous linear or polynomial fitting, is more aligned with the principle of letting the data speak for itself. In other words, we prefer to have the model follow the data, rather than the other way around (Benzécri, 1973).

The MSS approach can be conceptualized as an adaptive calibration process. Specifically, we seek to find the right balance between, on one hand, simply adopting a linear fit to the quantitative variable, and on the other hand, fully relying on the data and tracing the trajectory faithfully through each data point, regardless of the wiggliness of the fitting curve. Between the two extremes, the MSS approach endeavors to learn from the data by searching for a “sweet spot” through an adaptive process of optimization or partial pooling. This partial pooling mechanism has been applied to various scenarios in our previous work, including handling multiple testing (Chen et al., 2019a; Chen et al., 2019b; Chen et al., 2020a) and accounting for cross-trial variability (Chen et al., 2020b). The methodology elaborated here can be applied to any situation with a quantitative explanatory variable. Even though the demonstration data is of longitudinal nature, the modeling framework is not limited to longitudinal applications. In fact, it can be adopted for cross-sectional, population- or subject-level data as long as the range of the quantitative variable is wide enough to support a spline fit.

Here we introduce the program **3dMSS** that is publicly available as part of the AFNI suite (Cox, 1996) to the neuroimaging community for whole-brain voxel-wise analyses. In addition, we seek to

1. discuss the basic theory of MSS as a counterbalance between flexibility and stability;
2. lay out the schemes for population-level nonlinear analyses that involve experimental manipulation factors;
3. provide modeling accommodations to estimate and compare nonlinear effects;
4. demonstrate the modeling approach and visualization processes with a longitudinal dataset.

Throughout this article, regular italic letters in lower case (e.g., *β*) stand for scalar parameters or variables; boldfaced italic letters in lower (***b***) and upper (***X***) cases for column vectors and matrices, respectively.

## 2 Methods

### 2.1 Vanilla fitting: linearity

We start with the simplest scenario by laying down the data structure and model foundation. Suppose that we want to account for the effect of an explanatory quantitative variable *x* (e.g., age) on a response variable *y* at each spatial unit (e.g., voxel, surface node, region of interest, or whole-brain measure such as intracranial volume). Let {(*x_i_, y_i_*): *i* = 1, 2,… *m*} be the data from one subject with *i* indexing the *m* samples of the explanatory variable *x*. The distributional model can be formulated as follows,

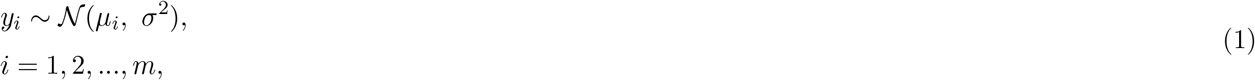

where *σ*^2^ is the variance for the likelihood (or prior) distribution (i.e., Gaussian) of the response variable *y*. The distribution mean *μ_i_* stands for the effect associated with the ith data point (*x_i_, y_i_*) that can be fitted as a linear relationship with the explanatory variable *x*,

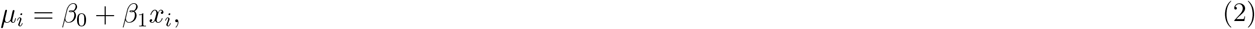

where *β*_0_ is the intercept while the slope *β*_1_ captures the marginal effect of *x*. The simple regression model can be reformulated in a concise vector-matrix form,

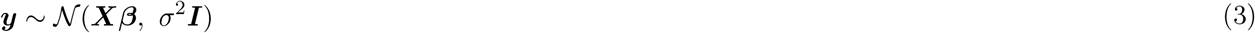

where vector ***y***_*m*×1_ = (*y*_1_,…, *y_m_*)^*T*^ represents the data for the response variable, ***X***_*m*×2_ = [**1**, (*x*_1_,…, *x_m_*)^*T*^] is a two-column model (or design) matrix, vector ***β***_2×1_ = (*β*_0_, *β*_1_)^*T*^ contains the two model parameters (intercept and slope) each of which is associated with a column in the model matrix ***X***. Under the conventional statistical framework, the regression or general linear model (GLM) in (3) assumes a Gaussian distribution; however, the exposition herein can be readily extended to the popular exponential family distributions such as binomial, Poisson and Gamma.

Linearity can capture long-range trends or overall associations between explanatory and response variables. The modeling strategy of straight-line fitting is frequently adopted in the literature because of its simplicity whenever a quantitative covariate is involved in, for example, a regression model solved through ordinary least squares (OLS). The slope directly reveals the marginal effect: a positive (or negative) slope indicates the amount of increase (or decrease) on average when the explanatory variable changes one unit. Similarly, correlation can reveal the overall strength of the relationship between two variables: a positive (or negative) correlation means that both variables generally move in tandem (or opposite direction). The graphic demonstration of linearity, as illustrated in the solid black line against the raw data of Fig. 1, is an omnipresent commodity in publications. For instance, it is fair to say that a substantial proportion of resting-state data analysis is based on linearity in the form of correlativity.

The information condensed in a linear model can be excessively crude. Whenever a local relationship or a short-range trend is of interest, linearity may fail to identify or may even distort the subtle relationship. For example, even though the linear fit in Fig. 1a provides a compact assessment of the general relationship, close examination shows substantially poor characterization across the left, middle and right segments of the *x* values: the predicted values of *y* are considerably larger than the raw data at both ends of the *x* values while it is the opposite problem in the middle range of *x*. When the relationship between two variables fluctuates or changes regularly or irregularly, the failure of the linearity assumption (Fig. 1b) may wreak havoc on data analysis and ultimately undermine efforts toward research reproducibility. It is this failure or distortion that necessitates more accurate modeling strategies than the default approach of linearity.

### 2.2 Modeling beyond linearity: spline interpolation

We intend to extend the linear model (2) by overcoming the limitations of polynomial fitting. Despite its unwieldy data fitting across the whole range, polynomials have several desirable properties such as differentiability, smoothness, numerical simplicity and computational efficiency. Instead of assuming one full polynomial function for the whole range of *x* values, we may divide the data domain into multiple segments and fit each segment with a separate polynomial and keep the desirable functional properties but avoid the pitfalls listed in the Introduction. To achieve a degree of adaptivity to a set of data under consideration, suppose that we fit the data {(*x_i_, y_i_*): *i* = 1,2,…, *m*} with a *smooth* function *f*(*x*) that replaces the linearity in the formulation (2),

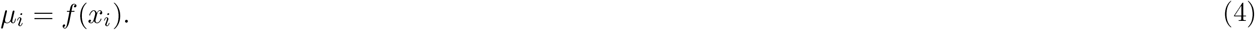

One solution is to fit *f*(*x*) through splines so that it does not have a specific predetermined form such as polynomial, exponential, logarithmic or trigonometric functions. Etymologically, the word *spline* refers to a flexible thin strip of wood or metal that can be used as a tool to draft different shapes of smooth curves. As illustrated by the line segment tool in Microsoft PowerPoint, various curves can be shaped with appropriate weights at various positions. Similarly, splines are smooth functions usually low-order polynomials with locations where adjacent splines meet each other as *knots*.

Two trade-offs are involved in the process - the spline type and the number of knots - both of which will have an impact on the fitness of the curves. The smoothness of a function is usually measured by its differentiability. A higher degree polynomial is more differentiable but computationally costly, while a lower order may result in jagged fitting around knots. Cubic polynomials are generally considered a well-balanced choice for their smooth smoothness at knots that are guaranteed through the the continuity up to the second order of derivatives. In addition, we adopt *natural* or restricted splines with their second and third derivatives set as 0 at the first and last knots so that the fitting is linear beyond the domain of the data at hand. As for the second trade-off, fitting with more knots may lead to overfitting, but smoothness and flexibility may suffer with a lower number of knots. For a specific set of knots, we build a set of basis functions so that any continuous function can be represented as a linear combination of basis functions.

We start with one type of natural cubic splines: cardinal basis functions. With *K* ordered knots *ξ*_0_ = *x*_1_ < *ξ*_1_ < … < *ξ*_*K*−1_ = *x_m_*, the constraints applied to natural cubic splines include: 1) *f*(*x*), *f′*(*x*) and *f″*(*x*) are continuous at the knots, and 2) *f″*(*x*) = *f′″*(*x*) = 0, when *x* = *ξ*_0_, *ξ*_*K*–1_. These constraints lead to *K* basis functions (and *K* free parameters) for *K* knots. With Δ*ξ*_*k*+1_ = *ξ*_*k*+1_ – *ξ_k_, β_k_* = *f*(*ξ_k_*), *γ_k_* = *f″*(*ξ_k_*), *k* = 0,1,…, *K* – 1, the smooth function in (4) can be parameterized (Wood, 2017) as,

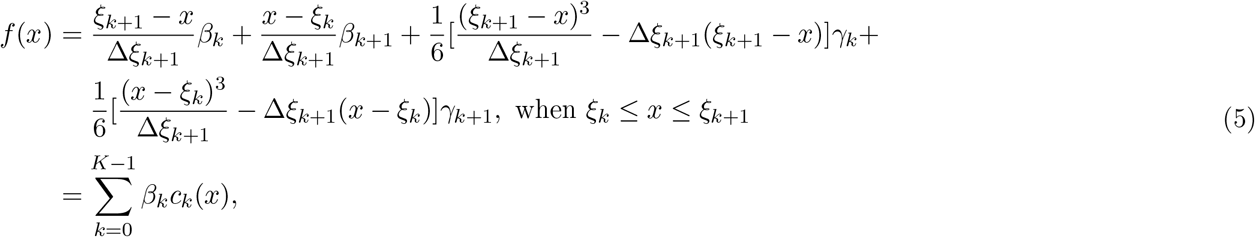

where {*c_k_*(*x*): *k* = 0,1,…, *K* – 1} are the cardinal cubic basis functions. One nice feature of the cardinal splines is that the *k*th basis function peaks with a value of 1 at the associated knot *ξ_k_* while having the value 0 at all other knots (*k* = 0, 1,…, *K* – 1) (see an example with *K* = 6 knots in Fig. 2a); thus, each coefficient *β_k_* can be directly interpreted as the fitted value at the knot *ξ_k_*. We can update the formulation (1) as

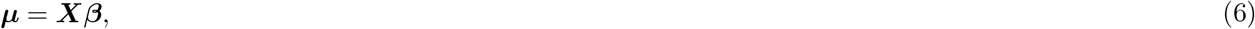

where ***μ*** = (*μ*_1_, *μ*_2_,…, *μ_m_*)^*T*^, ***β*** = (*β*_0_, *β*_1_,…, *β*_*K*–1_)^*T*^, and the model matrix ***X*** is of dimensions *m* × *K* with its (*i, k*)th element being *c_k_*(*x_i_*). The formulation (6) under the distribution (1) can be solved through, for example, OLS when *K* < *m*. However, overfitting cannot be effectively controlled under (6), and predictive accuracy may suffer when the model is applied to out-of-sample data. Thus, a balancing mechanism should be considered to counteract the danger of overfitting.

**Figure 2:**
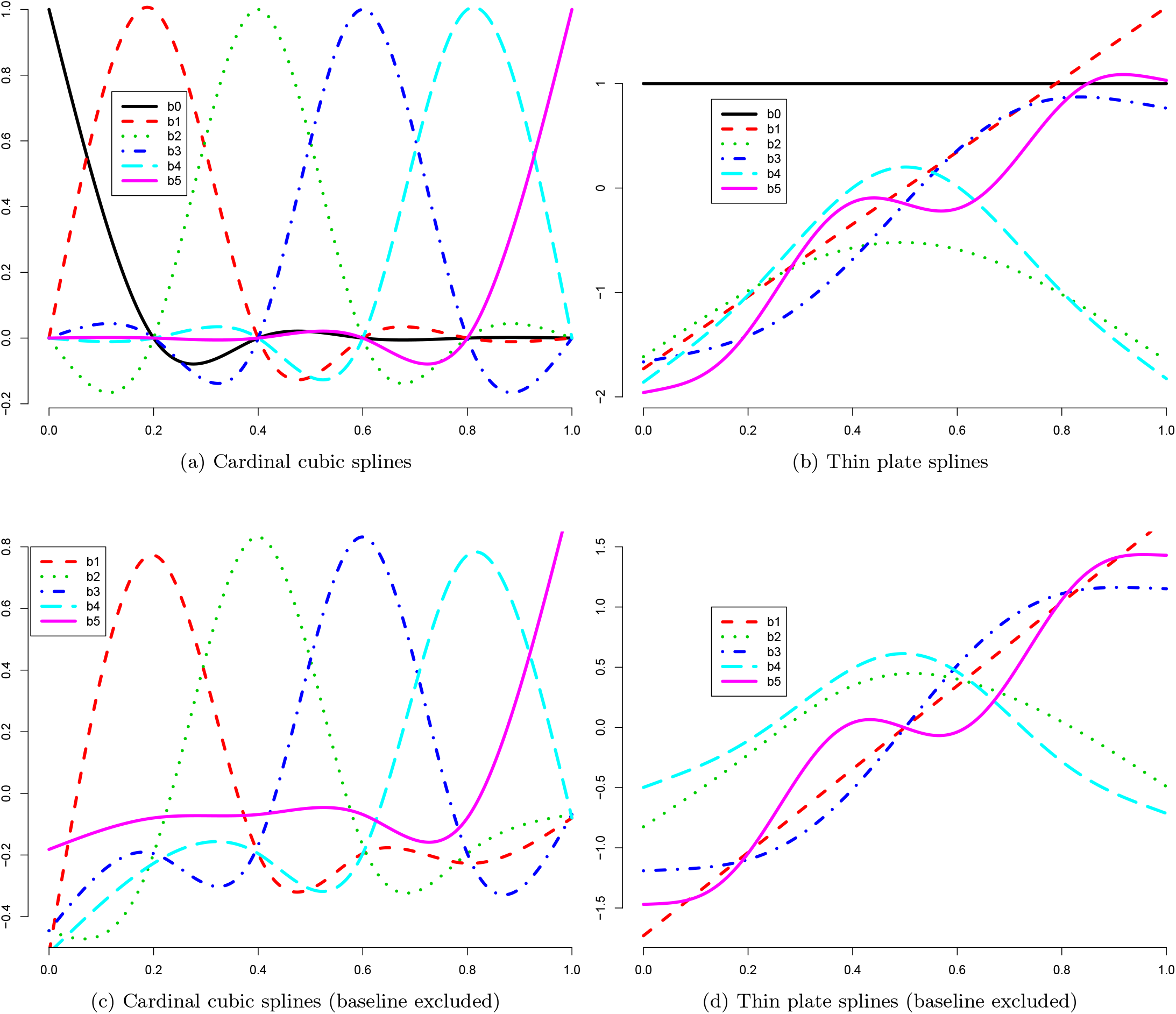
Basis functions *b*_0_(*x*), *b*_1_(*x*),…, *b*_*K*−1_(*x*) with *K* = 6. Two spline types are discussed here: cardinal cubic splines (a, c) and thin plate splines (b, d). Cubic splines are knot-based; with 6 knots at 0, 0.2, 0.4, 0.6, 0.8, and 1.0 within [0, 1], there are 6 cardinal basis functions (a). Each cardinal basis function (e.g., *b*_4_(*x*) in blue) peaks at the associated knot (*x* = 0.6) with a value of 1, and takes the value 0 at all other knots. Therefore, the fitted value at a knot corresponds to the weight for the associated basis function. In contrast, as shown in (b, d), each of the thin plate basis functions within [0, 1] is not knot-specific. As the baseline (intercept) is usually modeled separately in real practice, we only utilize *K* – 1 basis functions to maintain model identifiability (c, d).

### 2.3 Regularization through Bayesian multilevel modeling

Now we wish to resolve the second trade-off with respect to the number of knots. The challenge is to adaptively find a point of balance in the tug of war between two forces: too few knots may result in underfitting and rough curves, while too many would lead to overfitting or even an unidentifiability problem. The solution involves the second derivative of a function, which is related to the curvature or concavity of its curve: a positive second derivative corresponds to upwardly concave, and vice versa for a negative second derivative. Under the Bayesian framework, one technique to achieve adaptivity is partial pooling. When the potential risk of overfitting exists, we calibrate or leverage the peer effects (e.g., fitted values across multiple locations or knots) through a prior distribution. To differentially, instead of equally, calibrate those fitted values, we control the smoothness, roughness or curvature of spline function *f*(*x*) by tuning the integrated square of the second derivative that can be formulated (Wood, 2017) per (5),

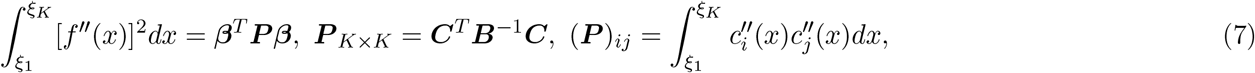

where the matrices ***B*** and ***C*** (defined in Appendix A) are composed of the (usually equal) distances Δ*ξ_k_* among two neighboring knots. With the notations ***Q*** = ***B***^−1/2^***C*** and ***b*** = ***Qβ***, we have ***Q***^*T*^***Q*** = ***P***. Now we formulate the spline model (6) under the Bayesian multilevel (BML) framework,

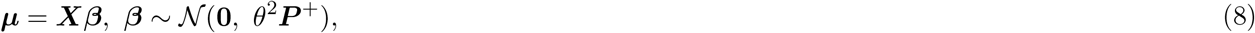

where ***P***^+^ is the pseudo inverse of the penalty matrix ***P*** and plays the role of regulation through the prior distribution 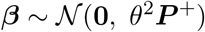. Similar to ***B*** and ***C***, the elements of ***P*** and ***Q*** are just the distances Δ*ξ_k_* among the neighboring knots. Thus, each element of the vector ***b*** is a weighted combination of the model coefficients *β_k_* that are associated with the cardinal basis functions *c_k_*(*x*). Essentially, the prior 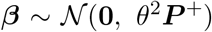 can be directly translated to a regularization on the basis weights 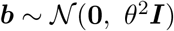.

The fitting adaptivity with a wide range of knots can be achieved through the Bayesian framework (8). Specifically, ***Xβ*** embodies the closeness of the fitting curve to the data, while ***b*** regulates the complexity of the curve through its distribution assumption. Specifically, the prior distribution of ***b*** plays the role of partial pooling as revealed by the two extreme scenarios. When the variance *θ*^2^ = ∞, *no pooling* is executed, the model fully interpolates the data and may result in rough or zigzagged fitting; on the other hand, *θ*^2^ = 0 means *complete pooling*, which corresponds to *f″*(*x*) = 0 or a linear fit. However, *θ*^2^, as a hyperparameter, is inferred from the data at hand; in other words, the amount of pooling between the two extremes is adaptively determined, and the specific number of knots becomes inconsequential as long as it surpasses some minimum threshold (e.g., 10).

One practical hurdle of adopting the BML framework (8) is the high computational cost. For whole-brain voxel-wise analyses, the computational burden depends on three factors: the number of subjects, the number of voxels and model complexity (e.g., number of knots, number of variables). Several features of Bayesian modeling are superior to the conventional approach, including straightforward results interpretation and presentation, easy incorporation of prior knowledge and flexible adaptation to data distributions. For whole-brain voxel-wise analysis, the computational challenge currently remains prohibitively costly, and we need alternative approaches to achieving the goal of regularization. Nevertheless, the BML model (8) provides a crucial platform for deriving uncertainties of predicted values in smoothing spline modeling.

### 2.4 Regularizing knots through LME and penalized least squares

One alternative to BML is to translate it to a linear mixed-effects (LME) counterpart. Denote ***β***^*^ = (*β*_0_, *β*_1_)^*T*^, and ***b***^*^ = ***Q***(*β*_2_,…, *β*_*K*–1_)^*T*^, and define two matrices 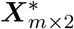 and 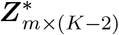 as the first two columns and the rest of the columns, respectively, of 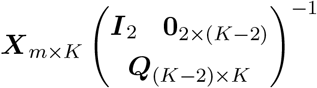. With the following reparameterization (Wood, 2017),

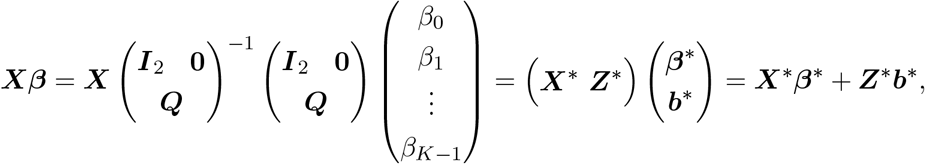

the BML model (8) is updated to an intuitive and familiar LME format,

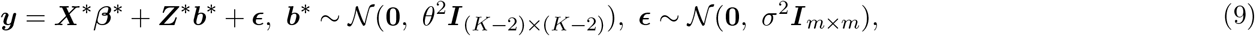

where the first two basis coefficients ***β***^*^ are conceptualized as fixed effects, and the remaining *K* – 2 components ***b***^*^ in ***b*** = ***Qβ***, as the combinations of all the basis coefficients ***β*** (including the first two), play the role of random effects that follow a Gaussian distribution, same as the prior distribution for ***b*** in the BML model (8). Under some scenarios, including the current context, the two frameworks of BML and LME are conceptually parallel with each other. However, the LME model (9) can be solved in much less computationally demanding ways, through, for example, optimizing the restricted maximum likelihood (REML).

Another alternative to BML is conventional ridge regression. Instead of minimizing (***y* – *Xβ***)^*T*^(***y* – *Xβ***) through OLS, we add a quadratic penalty term, the integrated square of the second derivative of *f*(*x*), to the regression model (6) and solve it through penalized least squares (PLS),

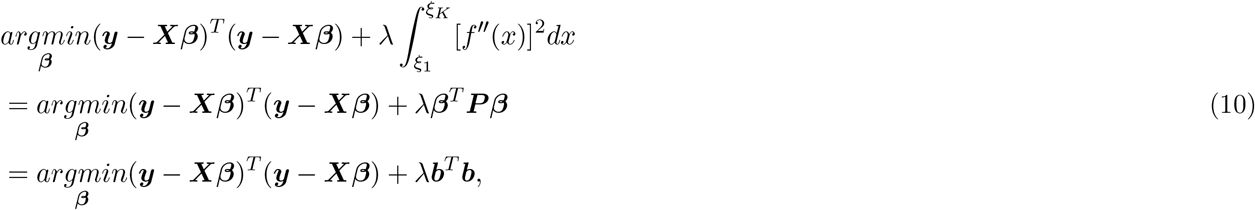

where λ ≥ 0 is the tuning or smoothing parameter. In contrast to the conventional ridge regression approach, here we do not equally penalize the basis weights ***β***, but instead apply the *L*^2^ regularization to ***b*** or various combinations of the basis weights ***β***. It is for this reason that ***P*** is referred to as the penalty matrix. Similarly, the intuitive interpretation of PLS is the counterbalance between the two forces of under- and over-fitting: λ = 0 corresponds to no penalization while λ = ∞ leads to full penalty (straight line). If λ is known, the PLS solution can be obtained through

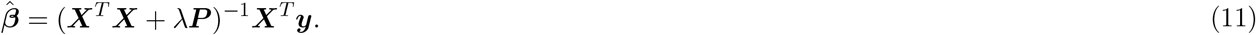

Similar to typical ridge regression, the smoothing parameter λ is usually determined through various crossvalidation methods or Akaike information criterion (Wood, 2017).

The three models - BML, LME and PLS - are essentially equivalent. In fact, their equivalence is represented by a correspondence between the penalty parameter λ and the relative magnitude of variance for the prior and random-effects distribution: 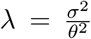. To put it another way, the prior distribution under BML can be alternatively expressed as 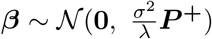, and the posterior distribution as 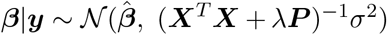. In addition, the LME formulation (9) can be directly treated as a BML model through the conceptualization of the distributional assumption of the random effects ***b*** as a prior. Each of the three modeling frameworks has unique pros and cons in terms of inference formulation, interpretation and computational consideration. In the current context, all three models are complimentary to each other in the development and applications of smoothing spline modeling.

### 2.5 Statistical inferences with smoothing spline modeling

To broaden the applicability in real practice and numerical implementations, a few modifications are warranted. First, to increase the flexibility in basis function selection, we consider a different type of smoothing basis, thin plate spline. Historically, thin plate splines were developed for two or more explanatory variables with the analogical concept of a thin sheet of metal that is resistant to bending (Duchon, 1977). Most types of basis functions (including the cardinal cubic set) for smoothing spline modeling are knot-based in the sense that a set of knots has to be specified before the basis functions are constructed as shown in the formulation (5), and the determination of knot number and their locations may add some extent of arbitrariness. By contrast, with thin plate smoothing splines we directly focus on the number of basis functions rather than the number of knots. Specifically, we start with as many basis functions as the unique sampled data points of the explanatory variable and then construct a specific number of basis functions by reducing the dimensionality through eigen-decomposition. In consequence, we adopt approximate thin plate splines and achieve a counterbalance between the effectiveness in smoothness regularization and computational cost (Wood, 2003; Wood, 2017). A set of thin plate splines with 6 basis functions within [0, 1] is illustrated in Fig. 2b.

Thin plate splines have some advantageous features that lead to intuitive interpretations. For example, the linear components of intercept and slope are part of the basis functions (*b*_0_(*x*) and *b*_1_(*x*) in Fig. 2b). Subsequently, we could parameterize the BML/LME formulation (9) in such a way that the two linearity components are directly associated with matrix ***X***\*. In other words, the two model coefficients *β*_1_ and *β*_2_ in ***β***\* are the intercept and slope of the linear components. This straight interpretation offers convenient model formulations and statistical inferences as elaborated below.

Research focus may vary in terms of the linear and nonlinear components in a model. One accommodation is to modify our basic formulation of smoothing splines in (4) by extracting the intercept or baseline from the smooth function *f*(*x*),

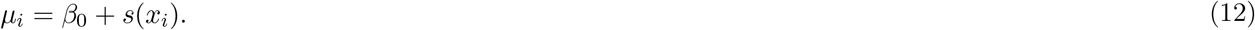

In this formulation the linear component remains part of the smooth function *s*(*x*). Specifically, column centering is required across the rows of matrix ***X***^*^ to achieve indentifiability under the intercept formulation (12); that is, we need to reduce the number of the basis functions from *K* to *K* – 1 (Fig. 2c,d). Another modification is to separate both the intercept and linear term from the smooth function *f*(*x*) in the formulation (4),

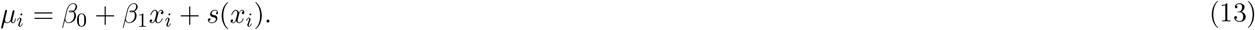

Under these reformulations, some adjustments are needed. Specifically, under the model formulation (9) shared by both BML and LME, reparameterization may be needed. Thus, as the baseline and linear basis functions are directly part of the thin plate basis functions, indentifiability can be accomplished through the reduction of the number of the thin plate basis functions from *K* to *K* – 2, for the formulations (13), by separating both the baseline and linear components (solid black and dashed red lines in Fig. 2b) from the thin plate basis functions.

Statistical inferences can also be performed separately for the linear components (or parametric coefficients, e.g. intercept *β*_0_ and slope *β*_1_) and the nonlinear components (or smooth terms) in *s*(*x*). The statistical evidence for each component in the linear terms is usually constructed through Wald test using the Bayesian covariance matrix for the coefficients. In contrast, a *χ*^2^-statistic can be adopted to assess the strength of the nonlinear terms using the uncertainty measures under the BML framework (Wood, 2013; Wood, 2017); in this case, the statistical evidence is assessed against the nonlinear term being s(x) = 0. Specifically, *s*(*x*) = 0 corresponds to an intercept (constant or horizontal line) in the formulation (12) or *f′*(*x*) =0 in the formulation (4). On the other hand, *s*(*x*) = 0 is associated with a linear relationship in the formulation (13) or *f″*(*x*) = 0 in the formulation (4). The smoothness of the fitted curve can be assessed by the effective degrees of freedom for the associated *χ*^2^-distribution.

In addition to statistical inferences, one is often interested in tracing the estimated trajectory from the data and visualizing the smooth relationship. Such an undertaking can be fulfilled through predictions based on the model. At specific values of the explanatory variable x that are not necessarily aligned with the sampled data *x_i_*, the expected responses are estimated through interpolation and extrapolation per the model generation mechanism. In this regard, the Bayesian framework offers another important mechanism in assessing the uncertainty (e.g., standard error) through the posterior covariance matrix of the basis coefficients *β_k_*.

### 2.6 Modeling extensions

So far, we have focused on the basic concepts of smoothing spline modeling. These models (e.g., (4), (5), (12) and (13)) can be adopted to capture the nonlinearity of between-subjects quantitative variables that do not vary within subject in the context of cross-sectional analyses (e.g. brain volume, cortex thickness, IQ, age). With just a few extensions, a wider range of applicability can be further achieved. For example, to be able to perform population-level data analysis with within-subject quantitative variables (e.g., age in a longitudinal study), we include subjects as a variable (factor) in the model, justifying the adoption of the adjective “multilevel” in the acronym MSS for multilevel smoothing splines. In the following three model variations of handling within-subject quantitative variables, we simultaneously intertwine two pooling processes: one pools the variations across subjects toward the population effects, while the other drags or penalizes the smoothness toward the straight lines.

#### 1) Varying-intercept MSS

Suppose that we want to account for the effect of a quantitative explanatory variable *x* (e.g., age) on a response variable *y* at each spatial unit (e.g., voxel, surface node, region of interest) from *n* subjects. Let (*x_ij_, y_ij_*) be the data from *j*th subject’s *i*th observation (*i* = 1, 2,…, *m_j_*; *j* = 1,2,…, *n*), where the number of observations, *m_j_*, may vary across the *n* subjects. A simple extended model for handling a dataset at the population level is to add a subject-specific intercept,

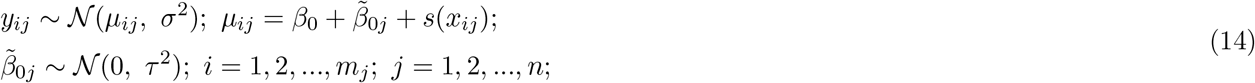

where *μ_ij_* stands for the effect associated subject *j* at the data point *x_i_, β*_0_ is the intercept or the population effect shared across all *n* subject, 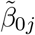 is the intercept deviation of subject *j* from the population intercept *β*_0_ and is assigned a prior Gaussian distribution with a variance *τ*^2^, and *σ*^2^ is the variance for the likelihood (or prior) distribution (Gaussian) of the response variable.

#### 2) Varying-intercept-and-slope MSS

Further extending the varying-intercept MSS model (14), we may allow both intercept and slope to vary across subjects,

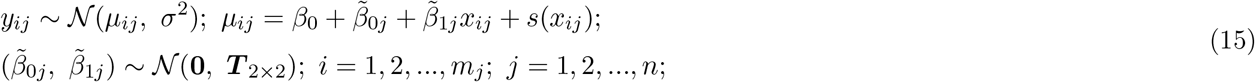

where 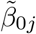 and 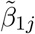 are the subject-specific intercept and slope, and their joint prior distribution is Gaussian with a 2 × 2 variance-covariance matrix ***T***. Notice that, with thin plate splines, the population-level slope effect is embedded in the smooth term *s*(*x_ij_*) in the formulation (15) with the corresponding basis function shown as the dashed red line in Fig. 2d. Nevertheless, one could extract the population slope effect as an explicit term if the assessment of nonlinearity is desirable.

#### 3) Varying-smooth-curve MSS

The most complex and most flexible extension is to allow each subject to have a unique smooth curve,

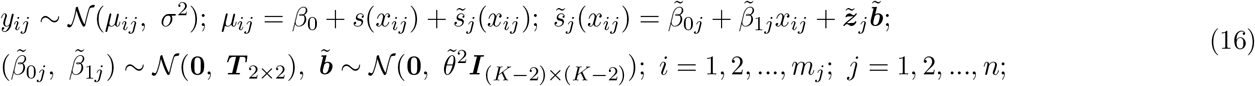

where 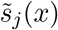 is the smooth fitting deviation of subject *j* from its population-level counterpart *s*(*x*), 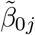 and 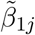 are the first two basis coefficients of intercept and linearity at the subject level that correspond to their counterparts ***β***\* in the BML/LME formulation (9) while ***T*** is the associated variance-covariance matrix for the joint prior distribution of those first basis coefficients, and 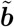 is the counterpart of ***b*** in the BML/LME formulation (9) while 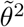 is the variance for the prior distribution of 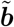.

All three population-level models, despite their varying complexity, can be conveniently parameterized, accommodated to and subsumed under the same BML/LME formulation,

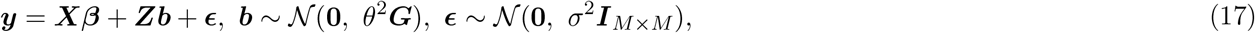

where ***G*** is a block diagonal matrix and 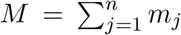. This unified model, as an extended version of its counterpart for a single subject (9), is achieved through matrix augmentation and integration across subjects, rendering an appealing platform for numerical algorithms. In fact, the previously elaborated numerical algorithms, statistical inferences and predictions can be readily applied to these three models.

### 2.7 MSS with more than one explanatory variable

Many data analysis scenarios, including neuroimaging analyses, may involve more than one explanatory variable. For example, an experiment may be designed with multiple groups of subjects or multiple conditions; in that case, a subject-grouping or within-subject factor has to be taken into consideration in the model formulation. Two modeling possibilities exist in terms of statistical inferences with MSS. The investigator may be interested in comparing the trajectories or trend of a quantitative explanatory variable across groups, conditions and various interactions. Alternatively, one could seek to focus on the interaction between the two experimental manipulation variables (e.g., groups and conditions) while adjusting the effect of a quantitative variable through smoothing splines instead of the typical approach of linear fitting.

Parallel to the conventional GLM and ANOVA platforms, a similar expansion can be formalized for MSS modeling. Here we utilize a concrete example to showcase the formulation process. Suppose that there are two experimental manipulation factors, forming a 2 × 2 factorial structure: one is a between-subject factor with two levels (patient and control) while the other a within-subject factor with two levels (positive and negative condition). We consider the data structure for two scenarios. One is to compare the trajectory or trend along a within-subject quantitative variable *x* (e.g., age) between the two groups, between the two conditions and their interaction, similar to the two main effects and the interaction in a conventional two-way ANOVA layout. In other words, the research hypothesis hinges on the comparisons among the smooth curves, not the comparisons of the average effects at a particular *x* value (e.g., mean). The other scenario is to perform ANOVA on the two factors while accounting for the effect of within-subject quantitative variable *x* as a confounder.

We formulate the ANOVA-analog MSS model through an old-fashioned dummy coding approach. Specifically, we choose a dummy coding method through which a factor of *L* levels is represented with *L* – 1 quantitative variables. With deviation coding, we set one level as baseline or reference that is coded as −1, and each of the other *L* – 1 levels is separately coded as 1. With two levels for both factors in this particular example, we formulate two quantitative variables, say d_1_ and d_2_, each of which is associated with one of the two factors and takes the values of −1 and 1. Then let *d*_3_ be the product of *d*_1_ and *d*_2_, representing the interaction between the two factors. Further suppose we adopt a varying-intercept MSS model as shown in (14) and use the indices *i, j* and *r* to track observations, subjects and the four factorial combinations between the two factors, respectively,

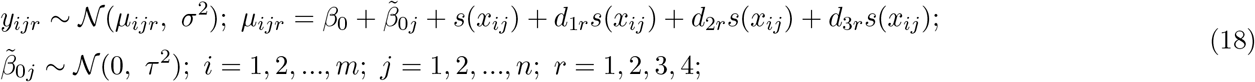

where *d*_1*r*_, *d*_2*r*_ and *d*_3*r*_ are the deviation coding values for each of the four factorial combinations. Under this formulation (18), *β*_0_ is the overall intercept at the population level accompanied with its subject-level intercept 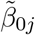; *s*(*x*) embodies the overall smooth trajectory; *d*_1*r*_*s*(*x_ij_*), *d*_2*r*_*s*(*x_ij_*) and *d*_3*r*_*s*(*x_ij_*) reveal the differences of the smooth trajectory between the two groups, between the two conditions, and the interaction (difference of difference) between the two factors, respectively. As discussed previously, the MSS formulation (18) can be parameterized and numerically solved under the BML, LME and PLS platforms. Furthermore, statistical assessments about the two main effects and the interaction regarding the smoothing trajectory can be completed as prescribed before, and so are the predictions for each of the four combinatorial scenarios.

The formulation for the ANOVA of the two factors with x as a confounding variable is more straightforward. With p and q coding the levels of the two factors, we decompose the effects as below,

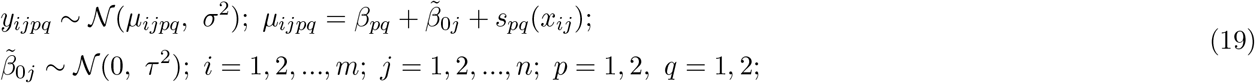

where 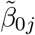 is the varying intercept with a prior Gaussian distribution 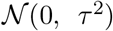, and *s_pq_*(*x_ij_*) is the smooth trajectory for the (*p, q*)th combination of the two factors. For meaningful interpretations of *β_pq_* and their various comparisons, the *x_ij_* values should be properly centered. Essentially, instead of a priori assuming a linear confounding effect of *x* as in the conventional treatment, here we replace a straight line fitting with more generic smooth functions *s_pq_*(*x*) in the formulation (19). While surely more computationally demanding, the MSS model (19) more accurately accounts for the data variability when nonlinearity is substantially present in the data.

One last modeling complexity is when there are two or more quantitative explanatory variables to be modeled through smoothing splines. Without loss of generality, we use two quantitative explanatory variables *x* and *z* as an example to briefly cover this extension, and simply expand, for example, the 1D formulation (12) with *s*(*x*) to,

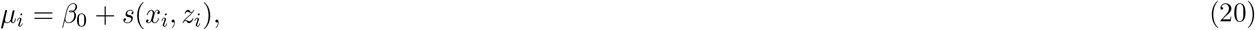

where *s*(*x, z*) is a bivariate smooth function. If *x* and *z* are isotropic in the sense that they are naturally on the same scale (e.g., 2D surface), a unit change in one marginal explanatory variable *x* corresponds to the same meaning as a unit change in the other marginal explanatory variable *z*; therefore, 2D thin plate basis functions can be directly adopted in this context to transform the formulation (20) to our familiar BML, LME and PLS model forms with only one prior distribution 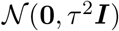, one tuning parameter *λ* and one penalty matrix as in the single quantitative variable case. On the other hand, when *x* and *z* are anisotropic in the sense that they are in different units (e.g., age and brain volume), we need two separate sets of basis functions; consequentially, we have two separate prior distributions, two different tuning parameters, and two separate penalty matrices in a tensor product form, one for each of the two explanatory variables. Nevertheless, despite the further complexity, the formulation (20) for the anisotropic case can, without exception, be turned into the same model platform as described previously (Wood, 2017).

### 2.8 MSS implementation for whole-brain voxel-wise analysis in neuroimaging

We offer an MSS program **3dMSS** that is publicly available as part of the AFNI suite for whole-brain voxelwise data analysis. Using the `R` (R Core Team, 2020) package **gamm4** (Wood, 2020), all the basis function options, including cardinal cubic splines, thin plate splines and tensor products are available. The modeling capabilities, as elaborated here, have been incorporated into **3dMSS** through a shell scripting interface (see a template in Appendix B). The respective model is applied at the voxel level through the conventional mass univariate approach; in other words, each voxel is fitted with unique smooth curves. All the explanatory variables and input data (in either text format for 1D and 2D or NIfTI format for 3D neuroimaging data) are fed through a table in the `R` long data format. Depending on the amount of data, spatial resolution, number of explanatory variables and model complexity, computational time may range from minutes to hours. Parallelization, making use of multiple CPUs and/or high-performance computing clusters, is recommended for demanding cases.

Statistical inferences and predictions are provided in the **3dMSS** output. The effect magnitude and the associated statistical evidence in *Z*-value are included for each of the linear components. The statistical evidence for each of the nonlinear components or their comparisons is assessed through a *χ*^2^-value; as the effective degrees of freedom vary across the spatial units, all the *χ*^2^-values are converted to have 2 degrees of freedom for the bookkeeping convenience of results storage. To make predictions for the trajectory of a quantitative explanatory variable, the user can specify the predictor values and factor levels through the long data format, and obtain prediction values and their uncertainty (i.e., standard errors) in the output. One can then interactively examine the trajectories and their comparisons, for example, at the voxel level through the graph window in the AFNI graphical user interface. The adjusted coefficient of determination 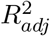 in the output can be utilized to assess the goodness of fit in model comparison. The conventional *R*^2^ can be directly interpreted as the proportion of variance explained by the model under, for example, GLM. The relative magnitude of the adjusted 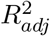 under MSS remains an important tool for model comparison, but the proportionality interpretation becomes somewhat opaque due to the presence of nonlinearity and the data hierarchy. In addition, the adjusted 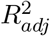 may occasionally become negative, indicating a model fit worse than a model based on an intercept alone.

## 3 MSS modeling of MRI brain data

To demonstrate the utility of the MSS approach, we applied the technique to a set of longitudinal VBM-style data cumulatively acquired from participants in the NIMH Intramural Longitudinal Study of the Endocrine and Neurobiological Events Accompanying Puberty (Reding, 2020), along with typically developing controls from The NIMH Longitudinal Study of Children with 7q11.23 Copy Number Variations (Gregory, 2019). This analysis was designed to assess the development of cortical asymmetries from childhood through adolescence. Previous cross-sectional adult studies examining asymmetry based on gray matter volume (Dorsaint-Pierre, 2006) or cortical thickness/area (Kong, 2018) have found hemispheric differences in planum temporale and/or Heschl’s Gyrus, which contain primary auditory cortex, but the developmental nature of these asymmetries remains to be fully characterized.

### 3.1 Experimental data

The MRI data information is as below. We acquired T1-based structural scans (MEMPRAGE, sagittal acquisition, 1-mm isotropic voxels, 256×256×176, TE/TR=1.828/10.556 ms, flip angle = 7°) on a GE 3T MR-750 scanner from 125 right-handed children (72 males, 53 females) in a longitudinal fashion, collecting 2-11 visits per participant over the age range of 7-20 years, for a total of 472 separate visits. Study procedures were approved by the NIH Combined Neuroscience IRB (NIH protocols 10M0112 and 11M0251). Participants were screened for eligibility based on history and physical examination by a clinician, by routine laboratory testing, and by a radiologist-reviewed MRI examination. Participants 18 or older and parents of minor participants provided written informed consent, and minor children provided assent. Three scans collected at each visit (each scan combined the four individual echos) were averaged and intensity-normalized, and gray matter was segmented in SPM12. Spatial normalization was performed within individuals and then on a group basis. Specifically, scans for all of an individual’s visits were spatially normalized to create a mid-time-point average image for that participant using SPM12’s longitudinal normalization tool (Ashburner, 2012). Advanced Normalization Tools (ANTS) software (Avants, 2009) was used to derive a symmetric group template based on structural images from 22 participant visits that were representative based on age and sex, including both the original and left-right reversed versions of each image. To ensure complete symmetry, the ANTS-based template was itself left-right reversed and averaged with the original. Following construction of the symmetric template, mid-time-point average images for each subject in the analysis were warped into this symmetric template space. Then, for each visit of each participant, ANTS was used to concatenate the deformation fields from the two spatial normalization stages (i.e., visit-to-subject-average, followed by subject-average-to-group-template), enabling the transformation to be performed with a single interpolation step. This transformation was used to warp gray matter images into the common symmetric group space, followed by smoothing at 8 mm and Jacobian modulation to account for local expansion or contraction based on the composite transform.

MSS modeling was adopted to explore asymmetries in developmental trajectories of cortical gray matter volume (GMV) on a voxelwise basis across the whole brain. For purposes of this analysis, original images from all participant visits were treated as belonging to one condition, while the corresponding left-right-flipped images were treated as belonging to subjects in a second (synthetic) condition. In other words, the asymmetry in gray matter developmental trajectory would be revealed through modeling the two conditions as two levels of a within-subject factor and contrasting the two conditions. From these data, three MSS models of voxelwise longitudinal gray matter trajectories across the brain were employed using **3dMSS** (scripts shown in Appendix B). The first model explored the asymmetry of developmental trajectories by contrasting the two conditions using the ANOVA platform of MSS (18) with the condition factor of two levels. The second model explicitly looked for the asymmetry of nonlinear trajectories after including both linear and nonlinear terms as formulated in the MSS model (13). As a basis for comparison of goodness-of-fit measures, the third model only accounted for the linear trajectory of age by adopting the MSS template (13) with the constraint *s*(*x_i_*) = 0. Predictions of gray matter volume trajectories were specified across the 7-20 year age range at specified half-year increments. With the input dataset consisting of 944 1-mm isotropic volumes, multi-threaded parallelization utilizing 16 threads within **3dMSS** yielded computation times of less than an hour for each of the output slices. Additional parallelization was made possible with a high performance computing cluster (the NIH HPC Biowulf cluster - http://hpc.nih.gov), such that each output slice was computed in parallel, allowing each of the two whole-brain voxelwise analyses to be completed in roughly one hour.

### 3.2 MSS modeling results

Our first model, based on the model (18), detected asymmetries of age-related trajectories in gray-matter volume with strong statistical evidence at several regions. The **3dMSS** output included the effect estimates and their statistical evidence in *χ*^2^-distribution for each parametric term in the specified model, as well as predictions of gray matter volume trajectories across the 7-20 year age range. The results shown in Fig. 3 illustrate findings of asymmetry in age trends of regional cortical gray matter volume as well as the statistical evidence through color-coded *p*-value. Specifically, the findings included a robust region of left-lateralized (left > right) asymmetry in Heschl’s gyrus, which is consistent with previous cross-sectional studies in the literature for adults (Dorsaint-Pierre, 2006; Kong, 2018). Examination of the spline-based trajectories of age by hemisphere shown in Fig. 3a offers some longitudinal insight regarding this asymmetry, indicating that this difference is likely present by the age of seven. Results also included (Fig. 3b) a region of similarly left-lateralized asymmetry in inferior temporal cortex, in which predicted trajectories showed an initial difference that diminished over the 7-20 age range modeled. Also, as illustrated in Fig. 3c, a small region was found within the anterior insula in which trajectories “crossed over” in the 14-15 year range. Although multiple regions demonstrated asymmetry with strong statistical evidence, the findings in these three regions were selected as a representative sampling of the types of differences between the original and flipped gray matter volumes in age trends that were discovered.

**Figure 3:**
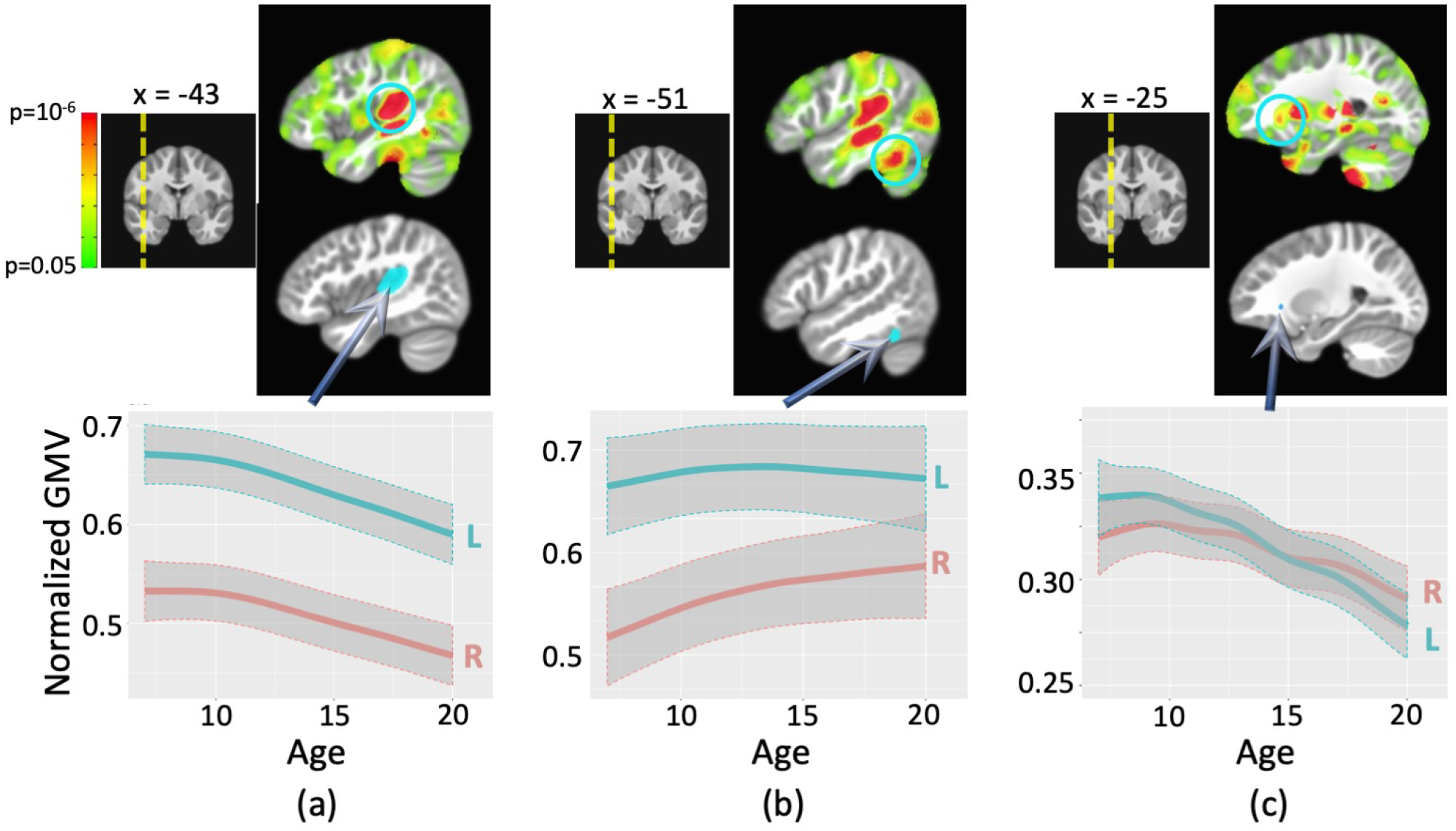
Statistical evidence of asymmetry in age trends of cortical gray matter volume (GMV) shown at voxelwise *p* < 0.05 (top) and three selected regions (highlighted in cyan) filtered at voxelwise *p* < 10^−5^ (below), based on smoothing spline modeling of trends with age. Trajectories are averaged across voxels within the three selected regions, and their uncertainty bands extend one standard error above and below the trajectories. Adjusted *R*^2^ values 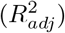 indicating goodness-of-fit were higher for the smoothing spline model than for a linear model (ranges for voxels within regions are shown below). (a) Heschl’s gyrus, demonstrating left-lateralized asymmetry with trajectories for left and right hemispheres that have roughly similar shape across the age range 7-20 (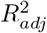 in (0.786, 0.997) for MSS vs. (−0.020, 0.353) for linearity); (b) Inferior temporal gyrus, also showing left-lateralized asymmetry, but with trajectories that reveal decreasing asymmetry over the same age range (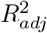 in (0.978, 0.998) for MSS vs. (0.065, 0.116) for linearity); (c) Anterior insula, where trajectories of right and left hemispheres can be seen to “cross over” (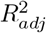 in (0.951, 0.983) for MSS vs. (−0.049, −0.022) for linearity).

Some subtle nonlinear trajectory differences were revealed from our second model. Fig. 4 illustrates findings based on the second model that specifically separated linear and nonlinear effects, showing the statistical evidence of regions for which asymmetry was expressed strongly in the nonlinear terms of the model. As the nonlinear components (with expected weaker statistical evidence) in the second model are embedded in the overall trajectory captured in the first model, Fig. 4 confirms that the regions associated with asymmetry based on nonlinear components were a subset of the regions revealed by the overall trajectory asymmetry (Fig. 3). Fig. 4a shows a right-lateralized region within the intraparietal sulcus (IPS) for which both hemispheres show gray matter reduction that increases in the 10-15 year range. Similarly right-lateralized regions within the temporo-parietal junction (TPJ shown in Fig. 4b) and and frontal pole (Fig. 4c) reveal an increased rate of reduction over approximately the same age range. The IPS region, thought to be a part of right-lateralized visuospatial processing systems (Kitada, 2006) and of a right-lateralized dorsal attention network (Corbetta, 2002), may undergo particularly aggressive specialization-based pruning during this time period. TPJ is likewise a recognized hub region, with evidence supporting its role in a right-lateralized ventral attention network (Thiebaut de Schotten, 2011) and thus may be similarly subject to accelerated pruning-based gray matter reduction within the same time frame.

**Figure 4:**
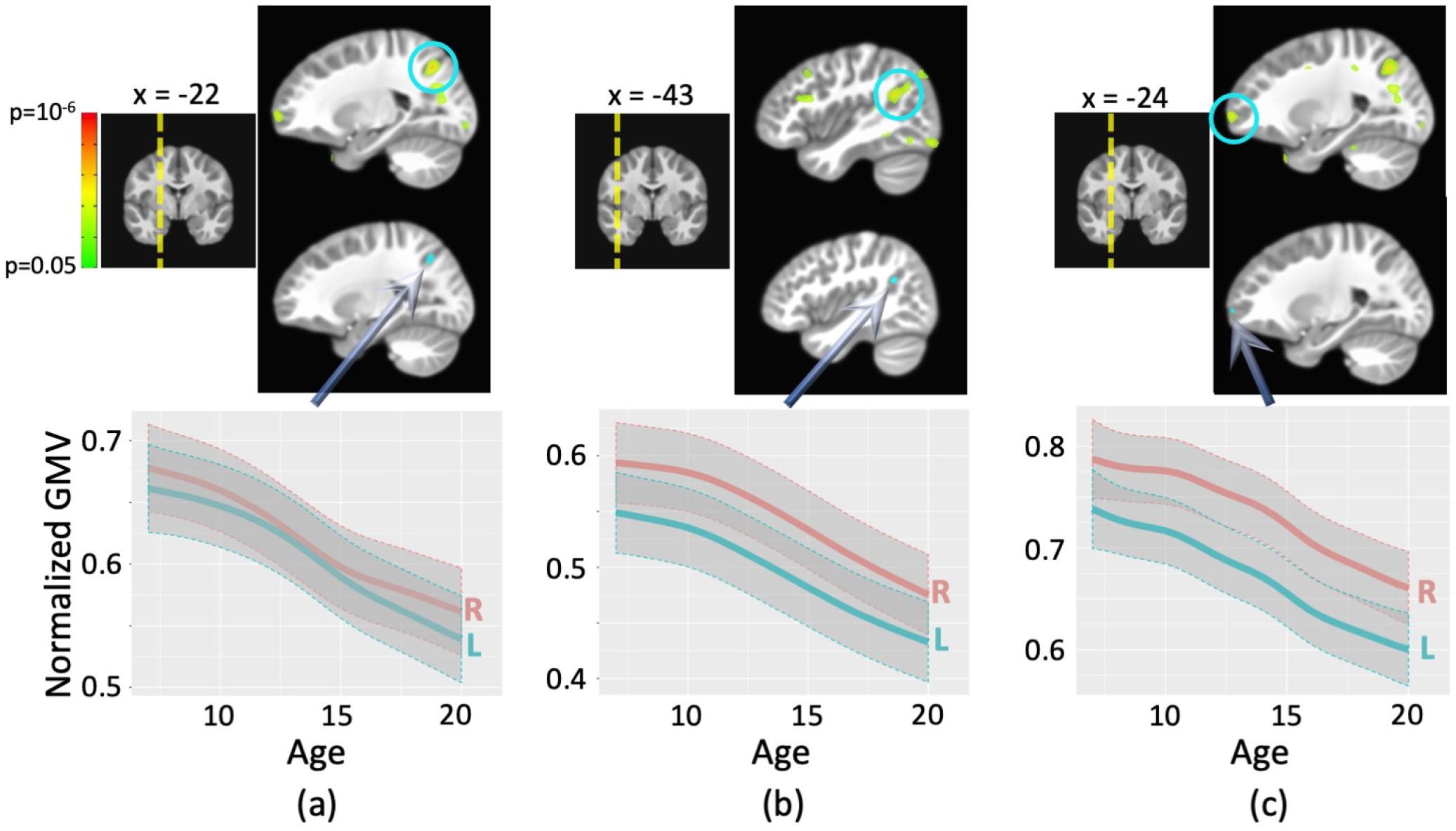
Statistical evidence of regions for which asymmetries were captured by a nonlinear trajectory (top, voxelwise threshold of p < 0.05) of cortical gray matter volume (GMV) and three selected regions (highlighted in cyan) (below, voxelwise p < 0.005). These regions were a subset of the general asymmetries shown in Fig. 3. The nonlinear trends are averaged within the three selected regions, and their uncertainty bands extend one standard error above and below the trajectories. (a) a right-lateralized region within the intraparietal sulcus where both hemispheres exhibit an increased rate of gray matter reduction in the 10-15 year range (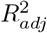 in (0.993, 0.996) for MSS vs. (0.001, 0.044) for linearity). (b) and (c) illustrate similarly right-lateralized regions within the temporo-parietal junction (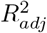 in (0.993, 0.998) for MSS vs. (−0.086, 0.018) for linearity) and frontal pole (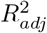 in (0.984, 0.991) for MSS vs. (0.051, 0.102) for linearity), respectively, with an increased rate of reduction over a similar age range.

## 4 Discussion

Linearity provides a convenient and irreplaceable framework for statistical models, and although it is appropriate for some experimental settings, its popularity can probably be largely ascribed to educational resource accessibility, familiarity and computational scalability. By virtue of effects assumed to be additive or accumulative, a linear model benefits from nice properties, numerical simplicity and intuitive interpretations. For example, OLS is computationally efficient in solving GLM; in addition, the Gaussian assumption guarantees the optimality of the effect estimate and the distributional properties for statistical inferences. It is no surprise that a large proportion of statistical data analysis is primarily performed under the umbrella of the staple methods such as regression, AN(C)OVA, GLM, LME and generalized linear model. These ubiquitous models are broadly covered in conventional statistical literature and are easy to use, visualize and conceptualize.

Neuroimaging data have historically been largely analyzed in a linear fashion. For example, the temporal dynamics of the BOLD signal in a brain region is constructed by convolving the stimuli with a linear time-invariant system and modeled as an additive effect in a time series regression model (e.g., Cohen, 1997). At the population level, a quantitative variable (e.g, age, cortex thickness), regardless of its being considered as an effect of interest or confound, is routinely modeled in a linear system.

### 4.1 Necessity of modeling nonlinearity

Nonlinear modeling broadens the capability and accuracy of data analysis. When the underlying mechanism is unknown or when a fundamental understanding of the root causes is deficient, the assumption of linearity as a first step may be helpful but not necessarily sufficient. Many relationships in child development, behavior, neurology, physiology and psychology are most likely nonlinear in nature and thus require appropriate analytic techniques. Additionally, nonlinear modelling can be an efficient way to explore complex phenomena when the specific relationships are initially opaque.

Among the wide variety of existing nonlinear modeling methods, smoothing spline modeling is well positioned as a tool for effectively capturing nonlinearity. At one end of the spectrum of methods, the analyst explicitly expresses the functional form with a fully parameterized model such as polynomials or logarithmic transformations. At the other extreme lie modern techniques that can arguably be described as “black box” approaches, including neural networks and their variations such as deep learning. Smoothing spline modeling positions itself somewhere in the middle of this spectrum.

Historically, the methodology of smoothing splines in a form suitable for functional data analysis first appeared 30 years ago, with the introduction of generalized additive models as an extension to generalized linear models in which the response variable can be linearly accounted for through unknown smooth functions of explanatory variables (Hastie and Tibshirani, 1990). It has been widely adopted in fields such as ecology, geology, meteorology and human health. Its applications in neuroimaging are gradually growing (Pomponio et al., 2020; Sørensena et al., 2020). The relatively easy adoption lies in the fact that once the number and type of basis functions are set, the model becomes linear in terms of the basis set, and the fitting proceeds as for conventional linear systems. Under flexible and relaxed assumptions regarding the actual relationship between the response and explanatory variables, smoothing spline modeling achieves improved fitting compared to conventional linear models, and strikes a balance between the goodness of fit, complexity and interpretability. A smoothing spline model is simply a GLM in which the explanatory variable partly depends linearly on some unknown smooth functions with the unknown nonlinearity transformed into a GLM through a set of smoothing splines. In parallel, an MSS model can be transformed into a BML or LME formulation, as elaborated in the Methods section. In fact, piecewise splines have been adopted to more accurately capture the hemodynamic response (HDR) shape variations; instead of a presumed HDR shape, tent basis functions or cubic splines can be utilized at the individual subject level (Chen et al., 2015) to achieve a higher modeling efficiency.

The characterization of nonlinearity through smooth splines hinges on the conventional concept of partial pooling. Without any prior knowledge regarding the specifics of the relationship, the only two prerequisites that play a significant role are the degree of smoothness and the prior distribution for the spline weights. The adoption of continuity up to second derivatives is partly for mathematical and pragmatical convenience (e.g., each knot is associated with a basis) and partly for satisfactory appearance (e.g., not too little or too much smoothing). The prior distribution for the basis coefficients amounts to applying a penalty against curve roughness to achieve a counterbalance in terms of the integrated square second derivatives. To put things in a different perspective, we resist the temptation to fit too close to the current data; rather, we hold a reasonable predictive accuracy as a desirable property when the fitted model is applied for interpolation and extrapolation. In the end, an overall compromise of the smoothness is reached through “borrowing strength” among the basis weights, a process to some extent comparable to the classical OLS in GLM, *L*^2^ regularization in ridge regression, and the calibration strategy in LME and BML.

### 4.2 Neuroimaging population-level analysis through MSS

Here, we present a multifaceted modeling platform that can be exchangeably formulated as Bayesian multilevel (BML) model, linear mixed-effects (LME) model, or ridge regression. This exchangeability among the three formulations allows us some conceptual convenience and flexibility in numerical algorithms as well as in statistical inferences. For example, fitting a smooth curve can be conceptualized as a regularization process among the basis coefficients under BML and LME; in contrast, smoothing spline modeling is analogous to a kind of “weighted” ridge regression in which the basis coefficients serve as penalty weights. The counterbalance between model complexity and data interpretability is accomplished through a calibration process of partial pooling with the roughness of the fitting curve penalized in a quadratic fashion. Just as in typical BML and LME frameworks, all the subjects under MSS are not required to have observations at the same *x* values (e.g., subjects scanned at different ages), and missing data are allowed as long as they are considered missing at random. Below we discuss four practical aspects of data analysis using smoothing splines.

#### 1) Choosing different types of smoothing splines

The basis set of thin plate splines is recommended unless the computational costs become unmanageable. Many basis types have been proposed over the years, but most of them either achieve roughly similar performance or are designed for niche usage (e.g., cyclic splines for periodicity). The two basis types discussed here are largely sufficient for most scenarios in neuroimaging: thin plate and cardinal cubic splines. The former has an intuitive aspect in the sense that its first two bases correspond to baseline and linear trend (black and red lines in Fig. 2b), offering the convenient flexibility to assess linearity versus nonlinearity as illustrated in the formulation (13) and in the second model adopted for our experiment dataset. On the other hand, the knot-based approach through cardinal cubic splines has a more direct interpretation with each basis coefficient corresponding to the estimated response magnitude. Practically, the thin plate spline basis is usually preferable by virtue of its slight edge in numerical performance (e.g., root-mean-square error) over other basis options including cardinal cubic splines. An approximate approach of thin plate splines starts with as many basis functions as the number of observations. To avoid wasteful computations and memory overhead, a reduction process of eigen-decomposition is adopted with a predetermined number of basis functions, *K*, and the first *K* components in the decomposed space are generated to retain most of the information through a low-rank basis set. This approximate basis set preserves much of the optimality of the conventional thin plate basis, but at considerably higher computational efficiency for large datasets. Another unique feature of thin plate splines is its baseline and linear components (solid black and dashed red lines in Fig. 2b). With the flexibility of the inclusion/exclusion of these two components, one can accommodate various inferential scenarios elaborated here including the model (13) that separates linearity from nonlinearity.

In some circumstances, cardinal cubic splines can be advantageous in terms of computation cost compared to its thin plate counterpart. Due to the massively univariate nature of typical neuroimaging analysis, the same model formulation is applied across the spatial units (voxel, region or surface node) repeatedly. Thus, the overhead setup time with thin plate splines through eigen-decomposition can be substantial with a large number of spatial units (e.g. voxel size of 1 *mm*^3^), a complex model or many subjects. In comparison, there is little model specification process associated with cardinal cubic basis functions because their knot-based expressions are explicitly available. While the results of the two types are virtually identical most of the time, the low setup cost of cardinal cubic splines may substantially cut the overall computational times.

#### 2) Determining the number of basis functions, *K*

The choice for the number of basis functions *K* is not as crucial and arbitrary as it appears to be. First of all, some minimum number of observations is required (e.g., at least 4 values per quantitative explanatory variable among all subjects) to fit a meaningful curve; otherwise, under- or over-smoothed fitting may occur. With adequate amount of data, small *K* leads to underfitting while large *K* may result in unnecessary computational cost. Therefore, a basic principle is that *K* should be sufficiently large (thereby avoiding bias from an oversimplified model) but small enough to retain computational affordability. In doing so, the specific *K* value would not be too critical with the following practical strategy. With more than 10 observations, *K* = 10 is likely a suitable choice for most situations to avoid incurring unwanted high computational burden unless the relationship is expected to have many twists and turns. On the other hand, with less than 10 observations, choose K to be close or equal to the number of observations.

#### 3) Tracking and comparing trajectories

Neuroimaging data analysis at the population level faces multiple challenges in modeling. First, the overwhelming amount of data is resource demanding on computational power. In addition, population-level neuroimaging analysis usually involves two or more cross-sectional dimensions. One hierarchical dimension is the cross-subject variability while another one is experimental manipulation variables such as groups (e.g., patients and controls) and tasks (e.g., positive, negative and neutral). With MSS modeling, yet another dimension is to handle the smoothness leveraging across basis functions. As a result, the analyst is not only motivated to track the trajectory under one particular condition for each group, but more likely interested in comparing trajectories across conditions or groups and exploring their potential interactions.

#### 4) Centering of a quantitative variable

Centering for a quantitative variable can be crucial for the interpretation of some effects. An explanatory variable is frequently categorized in practice into one of two varieties: either of interest or of no interest, with the latter typically labeled as a “covariate” in neuroimaging.^2^ For instance, whether and how such a covariate is centered is usually not revealed in the literature. As a practical issue, centering is trivial to apply and usually not discussed or emphasized in textbooks. In addition, any potential interactions of a variable with other explanatory variables in the model are likely not considered nor discussed in the literature. Nevertheless, centering can be important in proper interpretation and numerical stability, and in typical practice it may not be as straightforward to determine whether and when centering should be performed within (or across) groups (or conditions). On one hand, centering would not matter when the slope or the marginal effect of the quantitative variable is of interest. On the other hand, without centering, for example, the difference between two sexes would be interpreted when the quantitative variable (e.g. body weight) is adjusted to 0, which might not be necessarily meaningful. Furthermore, if the two groups intrinsically differ on average regarding the quantitative variable, centering becomes subtle or even pivotal as an effect of the group difference would be associated at zero, overall or group mean of the quantitative variable depending on how it is centered. For example, for the trajectory of a quantitative explanatory variable, centering is not necessary in **3dMSS**. On the other hand, when effects such as intercept (e.g., *β*_0_ in formulations (12) and (13)) and factor effects (e.g., *β_pq_* in the model (19)) are of interest, one should consider centering for a quantitative explanatory variable before providing the variable values as input for **3dMSS**.

### 4.3 Model formulation options

We believe that the MSS framework can work in parallel with traditional population-level analyses. Instead of a linearity assumption for a quantitative variable, a smooth function can be easily inserted into the conventional model structure. Through the introduction of smoothing splines for nonlinear modeling, we sketch out the modeling formulations for four analytical scenarios as routinely practiced in neuroimaging. First, one may pivot on either tracing the trajectory of the quantitative variable or treating it as a confound. In addition, given sufficient data, different levels of model complexity can be explored by considering cases of varying intercept, varying intercept and slope, as well as varying nonlinearity. Furthermore, through the same contrast coding schemes as traditionally employed for factors, we build up the model platforms that assist the analyst in comparing the smooth trajectories across conditions and groups as well as in investigating interaction effects. Lastly, it is possible to separate the linear effects from nonlinear and to make respective inferences through, for example, the model formulation (13).

It is worth noting that a quantitative explanatory variable in MSS modeling can be either within- or between-subject in nature. Without doubt, a within-subject design is usually preferable by virtue of the statistical efficiency and robustness. Using age as an example, we emphasize that a within-subject experiment with data collected across various ages from each subject (e.g., in a longitudinal study) would provide more accurate and efficient assessment about the age trajectory due to a lower within-subject variation than a cross-subject design. On the other hand, a within-subject design is not always pragmatic due to constraints in resources and time. Nevertheless, this does not imply that one could not model and trace the age trajectory in a between-subject study. In fact, with enough number of subjects, an MSS model with the data collected from a cross-sectional study would still allow the investigator to explore and trace the age trend provided that the quantitative variable spans a reasonably wide range, and can be applied to big datasets such as Human Connectome Project (HCP), the Alzheimer’s disease neuroimaging initiative (ADNI), the International Neuroimaging Data-sharing Initiative (FCP/INDI) and the UK Biobank. The modeling limitations with this kind of data are reflected in the inability to capture or account for within-subject variability in terms of varying intercepts, slopes or smooth curves across subjects, as shown in the model formulations (14), (15) and (16)

All modeling formulations elaborated here are directly available at the whole-brain voxel level via the program **3dMSS**. Through an experimental dataset, we demonstrated the scripting interface (Appendix B) and usability of **3dMSS**. To avoid potential bias introduced into the literature by selective reporting and to improve reproducibility, it is important to report full results and to focus on effect magnitude and its uncertainty, as illustrated in our demonstration of gray matter volume development trajectory results (Figs. 3 and 4). Modeling at the voxel level through **3dMSS** shares the same multiplicity issue as the typical mass univariate methodology in neuroimaging; therefore, one should take appropriate measure to adjust the voxel-level statistical evidence when making statistical inferences.

### 4.4 Advantages and limitations of MSS modeling

Model fit verification is important but largely absent in most neuroimaging analyses. In fact, it is almost an impossible undertaking in neuroimaging to visualize and verify a linear or polynomial fit due to the large amount of data and the same formulation imposed across the brain. For instance, the massively univariate approach cannot accommodate a situation where a quadratic fit may suit one region while a cubic or quartic function could be a better choice for another region. Besides modeling challenges, a practical hurdle is that, whether or not a research variable is of interest, the goodness of the linear fit is rarely examined or verified in neuroimaging. Essentially, the willingness to visualize, select and improve a model is hampered by the overwhelming amount of data and the inflexibility of the massively univariate approach.

Alternative and more flexible approaches exist, but they either focus on different aspects or are usually more computationally costly. For example, K-nearest neighbor regression maintains a counterbalance between bias and variance, but does not enforce any smoothness of the fitting relationship. On the other hand, smoothing spline modeling can be considered as a special case of functional data analysis (Wang et al., 2016) under which each sample element is considered to be a function. Another more adaptive framework is Gaussian process regression that assigns a probability distribution over an infinite number of possible functions and the mean function can be considered as the maximum a posteriori estimation. In fact, a smoothing spline model is equivalent to a special case of Gaussian process regression with a proper covariance function through kriging (Kimeldorf and Wahba, 1970; Bay et al,2016). However, these more generic methods typically pay a prohibitive computational cost with, for example, Gaussian process regression scaled as 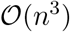 (Rasmussen and Williams, 2006), rendering them largely impractical especially when applied to neuroimaging data.

To summarize, smoothing spline modeling provides an elegant and compact analytical framework under which nonlinearity can be captured with minimal assumptions about the functional form that might best represents the underlying data. As long as the underlying relationship is more likely to be smooth rather than wiggly, the process of uncovering nonlinear effects through MSS becomes automatic and robust. In addition, much of the conventional model selection is now subsumed under the fitting formulation: the estimation of the basis coefficients is adaptively positioned on the basis of model simplicity, predictability and data fitness. Such adaptivity is especially important and appealing in neuroimaging, and can substantially alleviate issues of mis-fitting resulting from the inflexibility of conventional polynomial modeling when utilized in a massively univariate fashion. Aside from revealing potentially complex and interacting nonlinearity, the modeling framework also provides mechanisms for controlling, understanding and making inferences about those relationships. Sitting between model-driven and data-driven approaches, the methodology may serve as an exploratory tool before a bolder theory is proposed. The derived relationship is likely more robust and reliable than the alternatives determined via an assumption of linearity. The computational cost of MSS modeling for neuroimaging data is usually scalable and comparable to LME.

A few limitations of smoothing spline modeling are worth noting. With the linearity constraints (or zero curvature) levied upon the two end points of the observed data, the cardinal cubic splines approach may pay a price in poor predictability near the boundaries, especially when little information exists at the extremes of the range and beyond. Furthermore, the revealed relationship from the model does not necessarily provide insight into the relevant underlying physical/biological mechanism, nor can the model meaningfully make predictions beyond the range of the observed data. Lastly, if the sampled intervals of the quantitative explanatory variable are too large relative to the frequencies of the underlying mechanism, smoothing splines will likely result in over-smoothing or under-fitting; in this case, one cannot expect to estimate more than what the data could offer unless prior information or further constraints are available. For example, when the data are sparse or unevenly distributed within the modeled interval, the fitted trajectories, linear or nonlinear, might be excessively smooth; one should be more cautious toward the fitted trajectory and avoid solely relying on the statistical evidence.

## 5 Conclusions

Smoothing spline-based modeling extends conventional linearity-based modeling frameworks using quantitative explanatory variables. Under the assumption that the underlying relationship is likely to be smooth rather than rough, nonlinear effects can be effectively uncovered. The flexibility of MSS modeling may help uncover hidden patterns in the data without knowing a priori the specifics of an underlying functional relationship. The approach avoids the pitfalls of higher order polynomial fitting and can achieve a higher predictive accuracy. We hope that the MSS framework, with the associated program **3dMSS**, will contribute to improvements in neuroimaging data analysis.

## Acknowledgments

The research and writing of the paper were supported by the NIMH and NINDS Intramural Research Programs (ZICMH002888 for GC and RWC and NIMH ZIAMH002863 and ZIAMH002717 for KFB) of the NIH/HHS, USA, and a 2010 NIH Bench-to-Bedside award and a 2014 Brain and a Behavior Research Foundation Distinguished Investigator Award to KFB. The data for the NIMH Williams syndrome/typically developing cohort were obtained under protocol 10M0112/NCT01132885. The data for the puberty cohort were obtained under protocol 11M0251/NCT01434368, and we would like to thank Dr. Peter Schmidt, in collaboration with Dr. Karen Berman, for access to a preliminary dataset from their ongoing longitudinal study. We would also like to thank the participants in these protocols and their families, and to Gavin L. Simpson for his modeling advice. We thank the anonymous reviewers for their critical reading with thoughtful suggestions that helped improve, clarify and contextualize our manuscript. This work utilized the computational resources of the NIH HPC Biowulf cluster (http://hpc.nih.gov).

## Appendix A. Matrices *B* and *C* in (7) for smoothness regularization

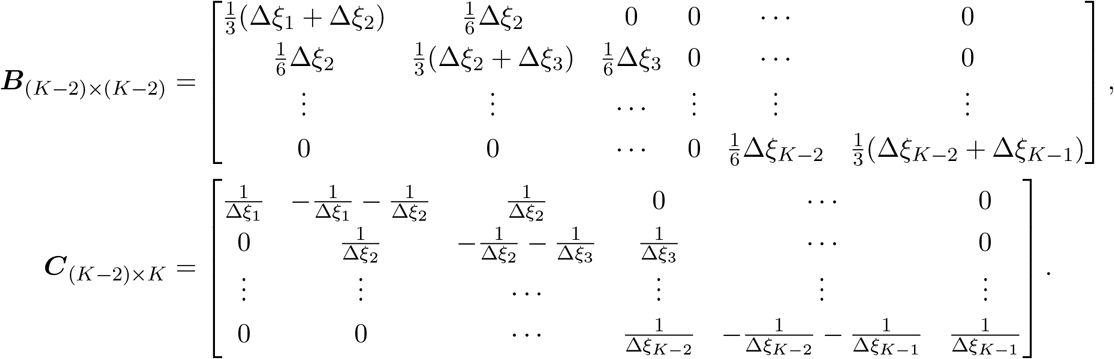

## Appendix B. Script and tables for running 3dMSS

The experimental dataset employed in the study contained two conditions of MRI brain volumes for each of the 125 subjects: one original and the other flipped. We intended to trace the age trajectories of gray matter volume from 7 to 20 years old and to assess the asymmetry by comparing the two conditions. The first MSS model compared the overall age trends and was specified with the AFNI program **3dMSS** as below:

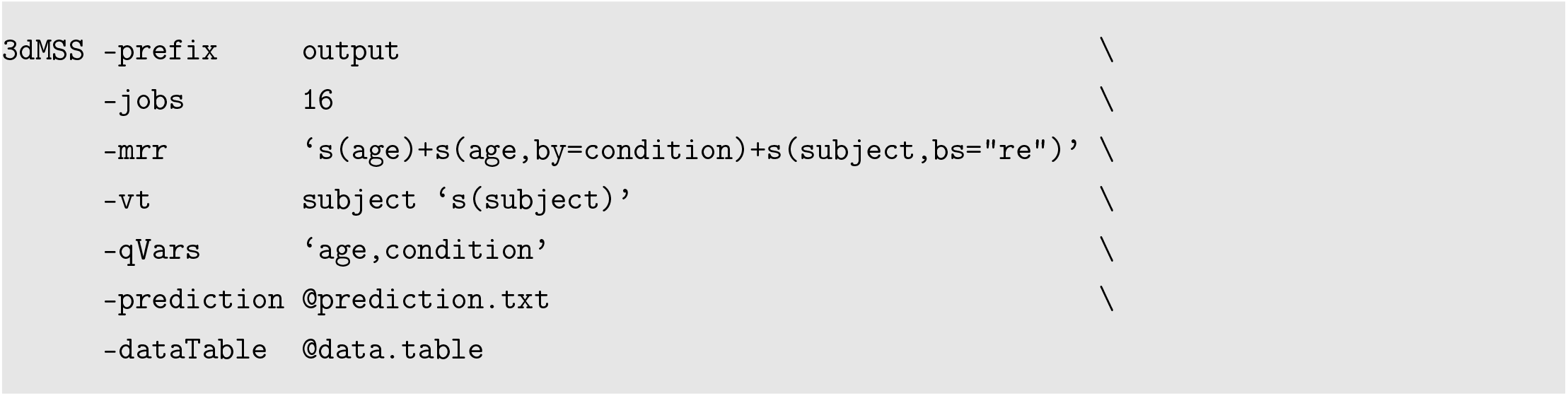

The output filename and number of CPUs for parallelization are specified through `-prefix` and `-jobs`, respectively. The expression *s*() in the formulation under model specification **-mrr** represents the smooth function, and the three terms `s(age), s(age,by=condition)` and `s(subject,bs=“re”)` code the overall age trajectory, the difference between the original and flipped conditions, and the cross-subject variability in intercept. The option `bs=“re”` indicates the random-effects variable (e.g., subject in this case). The number of thin plate spline bases was set to the default *K* = 10. The option `-vt` reveals the varying term (or random effects) in the model specification while `-qVars` identifies quantitative variables (`age` in this case plus dummy-coded `condition`). The last two specifiers `-prediction` and `-dataTable` list a table for prediction and input data information, respectively. A prediction table stored in the text file `prediction.txt` is of the following format:

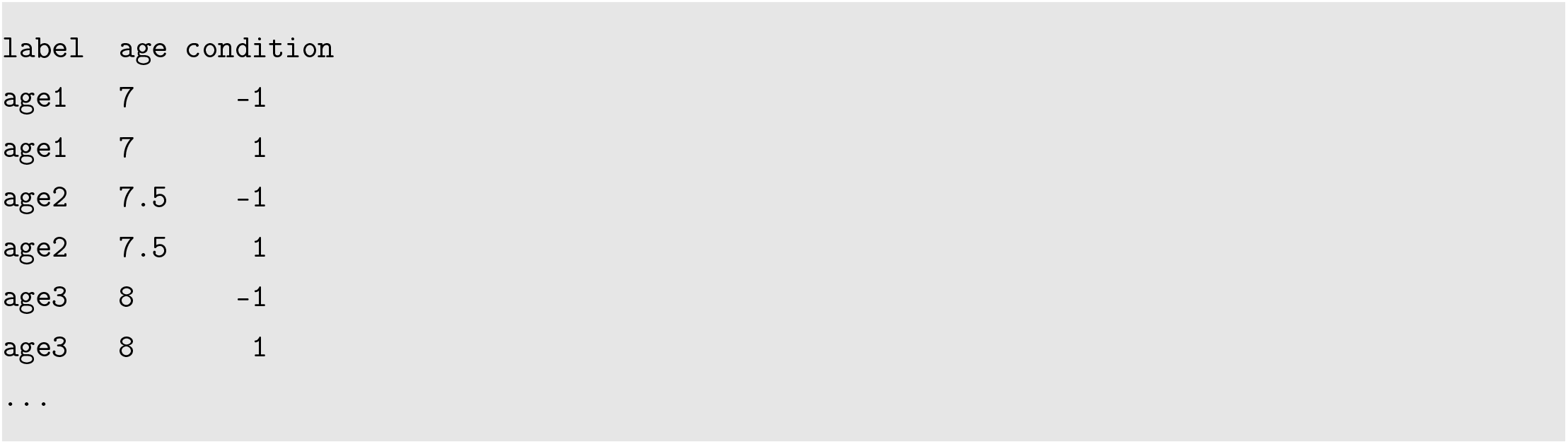

while the input data table has a structure as below:

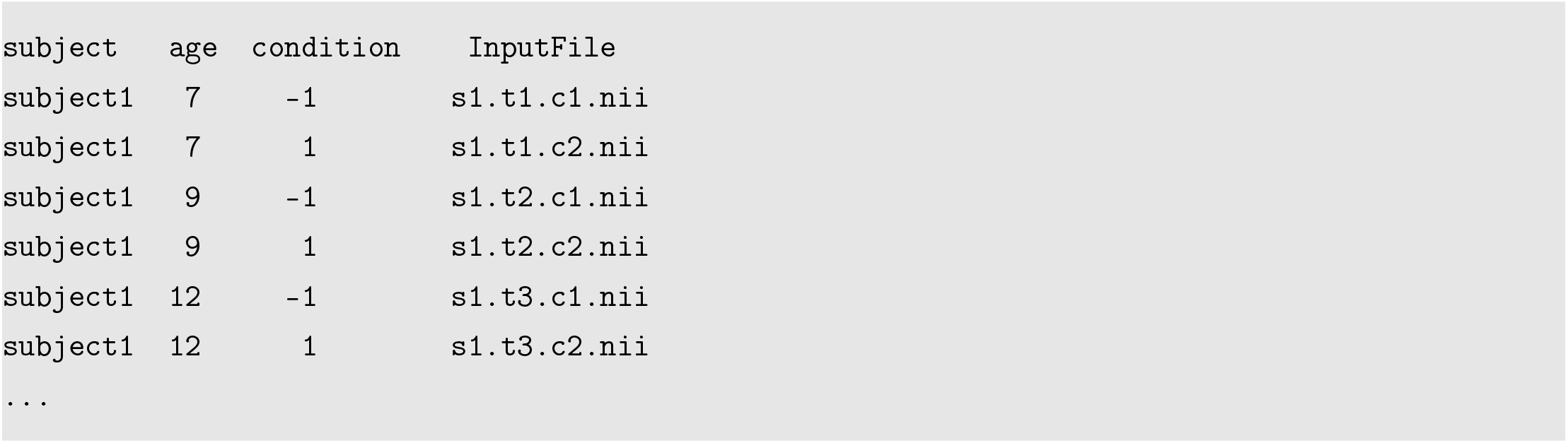

The second MSS model was constructed to capture and compare the nonlinear trajectories between the two conditions. The only difference from the first model was to replace the model specification by

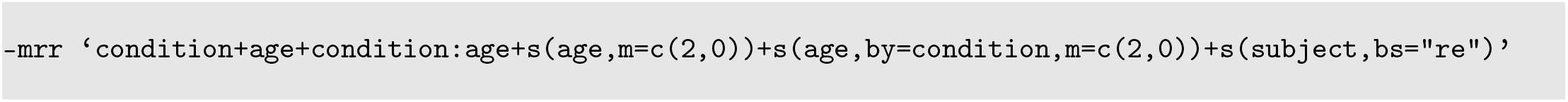

The linear effects were expressed in the terms `age` and `condition:age` while their nonlinear counterparts were modeled through `s(age,m=c(2,0))` and `s(age,by=condition,m=c(2,0))`. The option `m=c(2,0)` specifies a second order spline basis with 0th order penalty (ridge regression).

## Footnotes

1 In mathematics, a smooth function can be approximated by its Taylor series or polynomials in terms of the function’s derivatives at a single point.

2 The word *covariate* generally may have three different meanings in common usage. It is sometimes used to simply denote an explanatory variable that coexists with other explanatory variables in the same model. In the second popular usage, a covariate is a quantitative explanatory variable in the context of general linear model, while the third case is to mean an explanatory variable that is of no interest to the investigator. Likely due to software design, the word covariate is typically employed in neuroimaging to mean an explanatory variable, regardless of its nature (categorical or quantitative), that is of no interest to the investigator (e.g., age, sex, scanner, handedness, reaction time and intracranial volume). An implicit assumption in the widespread practice is that the effect of the explanatory variable, regardless of its strength, can be put behind the scenes as long as it is included in the model somehow. In some cases, the vague description and sometimes loose usage of the word covariate seem to serve as a protective umbrella so that the authors are immune to any request for the full revelation of modeling and result details.

